# Neural priority maps encode behavioral relevance independently across multiple attended locations

**DOI:** 10.64898/2026.04.29.721746

**Authors:** Amelia H. Harrison, Daniel D. Thayer, Thomas C. Sprague

## Abstract

Retinotopic cortical regions encode both the physical salience and behavioral relevance of visual stimuli, suggesting that these areas support neural “priority maps” of the visual field. These maps, which are instantiated as distributed neural codes across visual, parietal, and frontal areas, guide covert attention and overt motor plans. Previous work has characterized how these maps encode the task relevance of a single covertly-attended visual stimulus among multiple stimuli. However, it remains unknown how relevance-related modulations scale with the number of task-relevant locations. If modulations reflect limitations in attention-related enhancements of relevant neural populations, attending to more stimuli should reduce the magnitude of modulations. Instead, if relevance-related modulations reflect the selection of task-relevant populations by long-range inputs to optimize coding for further processing, each relevant stimulus location would be modulated by an equivalent amount, independent of other stimuli. We collected functional magnetic resonance imaging (fMRI) data while participants performed a demanding covert attention task requiring they monitor zero, one, or two peripheral task-relevant stimuli. Using a spatial inverted encoding model, we reconstructed neural priority maps from activation patterns across retinotopic cortex. Strikingly, relevance-related enhancement at each attended location was equivalent whether one or two locations were cued, with no evidence for attenuation when multiple stimuli were relevant. Moreover, task-related modulations were spatially focal and disjoint, localized to each cued stimulus location. These results suggest a model whereby behavioral relevance is encoded categorically in neural priority maps: BOLD signal from neural populations at each task-relevant location are equivalently and independent enhanced regardless of how many other locations are concurrently relevant. This modulatory profile reveals a ‘relevance map’ that signals which populations require selective augmentation to guide decision-making through the plethora of local neural computations associated with covert attention, with performance limitations arising at later processing stages rather than within early sensory representations.

## Introduction

Visually-guided behavior requires selecting relevant information from complex scenes for further processing. For example, when navigating a busy intersection, a driver must simultaneously monitor cars that might turn across their lane along with the traffic light ahead. To act efficiently, the visual system must adjust processing of relevant stimulus locations so that the most informative neural signals can be effectively communicated between regions to support goal-directed behavior.

Visual spatial attention serves this function by prioritizing sensory processing at task-relevant locations. Previous work in macaques and humans has demonstrated that attention modulates neural activity throughout early and mid-level visual cortex. In macaque electrophysiology studies, these modulations include increases in firing rate (Luck et al., 1997), burst rate (Anderson et al., 2011, 2013), neural sensitivity (Reynolds et al., 2000; Lee and Maunsell, 2010), reductions in activity fluctuations (Mitchell et al., 2007, 2009) and shared variability between neurons (Cohen and Maunsell, 2009), shifts in receptive-field position and size (Motter, 1993; Womelsdorf et al., 2006; Anton-Erxleben and Carrasco, 2013), and changes in inter-areal communication via altered synaptic efficacy and subspace alignment (Hembrook-Short et al., 2019; Ruff and Cohen, 2019). Human neuroimaging has revealed analogous effects in retinotopically organized regions across occipital, temporal, and parietal cortex. Attending to a spatial location results in enhanced BOLD fMRI signal (Kastner et al., 1998, 1999; Somers et al., 1999; McMains and Somers, 2004, 2005; Buracas and Boynton, 2007; Murray, 2008; Poltoratski et al., 2017; White et al., 2017; Itthipuripat et al., 2019a), changes in voxel-level measurements of receptive field properties (Sprague and Serences, 2013a; Kay et al., 2015; Sheremata and Silver, 2015; Tünçok et al., 2025), and selective enhancement of an attended stimulus location in multivariate population-level activation profiles (Sprague and Serences, 2013a; Kay et al., 2015; Vo et al., 2017; Sprague et al., 2018a).

Collectively, these mechanisms improve the quality and discriminability of neural representations at attended locations relative to unattended ones. A critical implication of this diversity of mechanisms is that each requires a “source” signal identifying *which* neural populations in sensory cortex encode behaviorally relevant information so that the appropriate "site" of modulation can be targeted for modulation. Consistent with this framework, a variety of causal and correlational studies in macaques and humans have identified changes in neural modulations in visual cortex as a function of spontaneous or stimulated activity in executive control regions in the prefrontal cortex and higher-order thalamus (Baldauf & Desimone, 2014; Ekstrom et al., 2008; Moore et al., 2003; Saalmann et al., 2012; reviewed in Squire et al., 2013).

While previous studies have evaluated how attention impacts activation profiles in early visual cortex when a single location is relevant, similar understanding of how attending to multiple locations impacts neural activity profiles is comparatively sparse. As fMRI BOLD signal is particularly sensitive to the combination of both long-range inputs to a region along with local neural activity (Logothetis et al., 2001; Logothetis and Wandell, 2004), experiments in which visual stimuli are held constant while relevance of multiple stimuli is independently manipulated can isolate the impact of top-down ‘source’ signals on the sensory ‘sites’ where stimulus processing occurs. Leveraging this approach, a previous study independently manipulating the luminance contrast of multiple stimuli under task conditions requiring focused attention to a single stimulus demonstrated a clear dissociation of stimulus-related and task-related signals across the cortex (Sprague et al., 2018a), as is typically observed in fMRI studies of spatial attention (Buracas and Boynton, 2007; Murray, 2008; Itthipuripat et al., 2019a).

These results have been used to support the contribution of different brain regions to various aspects of a neural ‘priority map’ that indexes the relative importance of each visual field location as a function of both its task relevance, independent of stimulus properties, and its stimulus salience, independent of task relevance (Itti and Koch, 2001; Fecteau and Munoz, 2006; Serences and Yantis, 2006; Bisley and Goldberg, 2010). While previous work has provided extensive evidence for ‘salience maps’ (where activity is driven by salient stimuli; Bogler et al., 2011; Burrows & Moore, 2009; Chen et al., 2020; Li & Wong, 2024; Liu et al., 2025), evidence for a salience-independent ‘relevance map’, where activity is driven only by task demands, remains sparse. Here, we test the hypothesis that top-down ‘source’ signals are reflected in a relevance map which highlights differences in priority map activation due only to changes in task demands. This map would reflect the spatial pattern of top-down inputs that instantiate the myriad mechanisms of neural response modulation associated with covert spatial attention that enable improved information processing and transmission.

To test this hypothesis, we characterized the spatial profile of these relevance-related modulations during task conditions requiring covert monitoring of multiple stimulus locations simultaneously. We independently manipulated the relevance of two identical stimuli presented at random positions on each trial while acquiring fMRI data across retinotopic cortex to visualize and quantify task-related modulations of neural priority maps. If a relevance map drives local changes in neural response properties to enhance information processing regardless of how many locations are concurrently relevant, relevance-related activation should be equivalent and independent at each cued location in a neural priority map. Instead, if the ability of the visual system to select and modulate relevant signals in neural priority maps is itself a major limitation in visual processing, the degree of relevance-related modulation should inversely scale with the number of relevant locations.

To preview our results, we found that activation at cued locations was constant as the number of task-relevant items was manipulated. This pattern held steady across retinotopic cortex. These findings suggest that relevance is encoded categorically and independently across locations throughout the visual processing hierarchy, such that multiple locations can be designated as relevant for enhanced processing and readout without apparent constraints.

## Materials and Methods

### Experimental Design

10 subjects recruited from the University of California, Santa Barbara (UCSB) community participated in the study (8 female, 21-33 years old), including two authors (AH and DT). All subjects had participated in other studies in the lab previously. All subjects reported normal or corrected-to-normal vision and did not report neurological conditions. Procedures were approved by the UCSB Human Subjects Committee and by the US Army Research Lab Human Research Protection Office. All subjects gave written informed consent before participating and were compensated for their time ($20/h for scanning sessions and behavioral sessions).

Each participant performed a 1-h training session before scanning, during which they were familiarized with all tasks performed inside the scanner. We also used this session to establish initial behavioral performance thresholds by manipulating task difficulty across behavioral blocks (see below).

We scanned participants for two 2-h main task scanning sessions. Each comprised three mapping task runs, used to independently estimate encoding models for each voxel (see **Fig. 3B**), and four selective attention task runs (broken into two sub-runs each, see below; used to characterize neural priority maps during focused and divided attention tasks). All participants also underwent additional anatomical and retinotopic mapping scanning sessions to independently identify regions of interest (ROIs; see *Region of interest definition;* **Fig. 3A**).

We presented stimuli using the Psychophysics toolbox (Brainard, 1997; Pelli, 1997) for MATLAB (The MathWorks, Natick, MA). Visual stimuli were displayed ∼149.5 cm from the participant’s eyes at the head of the scanner bore using a BOLDscreen 32 LCD display designed for the fMRI environment during the scanning session. In the behavioral familiarization session, we presented stimuli on a contrast-linearized LCD monitor (2,560×1,440, 60 Hz) 62 cm from participants, who were seated in a dimmed room and positioned using a chin rest. For all sessions and tasks (main tasks, retinotopy, and mapping task), we presented stimuli on a neutral gray circular aperture (7.72° radius), surrounded by black (only aperture shown in **Fig. 2**). Eye position was monitored throughout the experiment using an Eyelink 1000 eye tracker (SR Research) at 500 Hz.

### Selective attention task

We instructed participants to attend to random line stimuli (RLS). On each trial, the presented RLS would cohere into a “spiral” form, and participants responded with one of two button presses indicating which direction of spiral they detected in the cued RLS, or in the divided attention task, the first detected spiral (**Fig. 2A**). The RLS were presented at either 20%, 40%, or 80% contrast. We modeled this stimulus after that used in previous work (Sprague et al., 2018), in which the stimulus was designed to minimize the allocation of non-spatial feature-based attention, as well as to minimize the influence of potential radial biases in orientation preference in visual cortex as a function of preferred polar angle (Freeman and Simoncelli, 2011; Freeman et al., 2013).

Each trial began with a 750-ms spatial cue at fixation indicating the location(s) of the task-relevant stimulus/stimuli. The red arm(s) of the cue indicated with 100% validity whether each peripheral stimulus location was relevant on that trial. When no arms were red, the participant attended to the fixation point (Attend fix) (**Fig. 2A**). This was followed by either appearance of one or two RLS. For the attend-one condition, one RLS appeared at the cued location, and a second RLS appeared at another non-cued location (unless the non-target stimulus was not presented, see below). For the attend-two condition, both RLS always appeared at the two cued locations. Both stimuli remained onscreen for 3,000 ms, during which time a 1,000-ms spiral target appeared independently and instantaneously at each stimulus position. The target onset was randomly chosen on each trial for each stimulus (attended and unattended) from a uniform distribution spanning 500 –1,750 ms.

Both stimuli always appeared along an invisible iso-eccentric ring 4.22° from fixation. On each trial, the two stimuli appeared either 72° or 144° polar angle apart (4.96° and 8.03° distance between centers, respectively; **Fig. 2B; Fig. S1**). We randomly rotated the stimulus array on each trial around fixation, so the positions of the stimuli on each trial were entirely unique. Each trial was separated by an intertrial interval (ITI) drawn from a uniform distribution spanning 5.25– 9 s at 0.75-s steps, resulting in an average trial duration of ∼11.25 s.

Both stimuli flickered in-phase at 15 Hz (2 frames on, 2 frames off, monitor refresh rate of 60 Hz). Each stimulus consisted of 85 light and 85 dark lines, each 0.3° long and 0.035° thick, that were replotted during each flicker period centered at random coordinates drawn from a uniform disk 1.27° in radius. On flicker periods with no targets, the orientation of each line was drawn from a uniform distribution. On flicker periods with targets, the orientation of a random proportion of lines (defined by the target coherence) was either incremented (i.e., adjusted counterclockwise, “Type 1 spiral”) or decremented (i.e., adjusted clockwise, “Type 2 spiral”) from the Cartesian angle of each line relative to stimulus center (see **Fig. 2A**). Participants responded with a left button press to report a Type 1 spiral target or a right button press to indicate a Type 2 spiral target.

Stimulus contrast varied across trials (20%, 40%, 80%). Here, we focused on trials with two stimuli presented at matched contrast. However, in the acquired dataset, participants performed this task with a full counterbalanced set of stimulus contrasts (all combinations of 20%/40%/80% for one stimulus, and 0%/20%/40%/80% for the other stimulus, see **Fig. S1A**). Because we were primarily focused on the impact of instructed task relevance on activation profiles of neural priority maps, for this report we concentrated our analyses on trials with matched contrast across the two stimuli. For the full dataset, we fully counterbalanced the contrasts of each stimulus (above), task condition (Attend fixation, Attend 1, Attend 2), and separation distance (72° or 144° polar angle; **Fig. S1B**). Accordingly, to sample a single repetition of all trial types, we acquired 60 trials, broken up into 2 sub-runs, each lasting 345 s (30 trials, ∼11.25 s each, 2.25-s blank screen at beginning of scan, 5.25-s blank screen at end of each scan). Participants performed 8 full runs (16 total sub-runs), resulting in a full data set of 240 trials per participant. To keep performance below ceiling and at an approximately fixed level (at ∼80%), during scanning we adjusted the coherence of targets (defined as the percentage of lines forming a spiral target on each flicker cycle; **Fig. 2A**) independently for 20%, 40%, and 80% contrast stimuli in each attention condition before the start of each full run (i.e., 2 sub-runs). Initial coherence for each contrast and attention condition was calibrated during the pre-scanning behavioral familiarization session by manually adjusting values between runs.

When cued to attend to fixation, participants attended a flashing cross within the fixation circle and ignored any other stimuli presented throughout the trial (task adapted from Thayer & Sprague, 2023). Participants monitored the fixation cross throughout the trial for any increase in length in either the vertical or horizontal bar of the cross and responded to changes with a button press (left button for a horizontal target, right button for a vertical target). The vertical and horizontal lines of the fixation cross had a length of 0.13° of visual angle and flickered at 3 Hz (10 frames on, 10 frames off at 60 Hz). When a change was detected, participants reported which line increased in length (horizontal or vertical). To ensure equal task difficulty across attentional conditions, we adjusted the difficulty of the fixation task between runs by altering the degree of size change for vertical/horizontal lines based on behavioral accuracy. Although participants only responded to the cross when cued to fixation, the cross with associated targets was present through all trials.

### Spatial mapping task

We also acquired 6 runs per participant of a spatial mapping task to independently estimate a spatial encoding model for each voxel, following previous studies (Sprague et al., 2016, 2018; Sprague & Serences, 2013; Thayer & Sprague, 2023, 2025). On each trial, a flickering checkerboard (1.27° radius, 70% contrast, 6-Hz full-field flicker) appeared at a position drawn from a triangular grid of 37 possible positions, with a random uniform circular jitter (0.42° radius) added on each trial. The base grid separated adjacent stimuli by 1.9° and extended 5.69° from fixation (3 steps); combined with the jitter and stimulus radius, this yielded a visual field of view of 7.38° from fixation for the spatial encoding model. We rotated the base grid position on each run to increase spatial sampling density (Sprague et al., 2016), such that every mapping trial was unique. On trials in which the checkerboard overlapped fixation, we drew a small aperture around the fixation point (0.67° diameter). Each trial consisted of a 3,000-ms stimulus presentation followed by an either 6,000- or 8,250-ms ITI. Participants monitored for rare contrast changes (6 of 43 trials per run, 14%), evenly split between increments and decrements; on target trials, the checkerboard dimmed or brightened for 500 ms, beginning at least 500 ms after stimulus onset and ending at least 500 ms before offset. Task difficulty was adjusted between runs by varying the magnitude of the contrast change. All target-present trials were discarded prior to encoding model estimation. Each run totaled 432 s, including a 3-s blank period at the beginning and a 10.5-s blank period at the end.

### MRI acquisition

We scanned all participants on a 3T research-dedicated Siemens Prisma scanner at the UCSB Brain Imaging Center. fMRI scans for experimental, model estimation, retinotopic mapping, were acquired using the CMRR MultiBand Accelerated EPI pulse sequences. We acquired all images with the Siemens 64 channel head/neck coil with all elements enabled. We acquired both T1- and T2-weighted anatomical scans using the Siemens product MPRAGE and Turbo Spin-Echo sequences (both 3D) with 0.8 mm isotropic voxels, 256 x 240 mm slice FOV, and TE/TR of 2.24/2400 ms (T1w) and 564/3200 ms (T2w). We collected 192 and 224 slices for the T1w and T2w, respectively. We acquired three T1 images, which were aligned and averaged to improve signal-to-noise ratio.

For all functional scans, we used a Multiband (MB) 2D GE-EPI scanning sequence with MB factor of 4, acquiring 44 2.5 mm interleaved slices with no gap, isotropic voxel size 2.5 mm and TE/TR: 30/750ms, and P-to-A phase encoded direction to measure BOLD contrast images. We measured field inhomogeneities by acquiring spin echo images with normal and reversed phase encoding (3 volumes each), using a 2D SE-EPI with readout matching that of the GE-EPI and the same number of slices, no slice acceleration, TE/TR: 45.6/3537ms.

### MRI preprocessing

Our approach for preprocessing, adapted from our previous studies (Thayer and Sprague, 2023, 2025), was to coregister all functional images to each participant’s native anatomic space. First, we used all intensity-normalized high-resolution anatomic scans (3 T1 images and 1 T2 image for each participant) as input to the “hi-res” mode of Freesurfer’s recon-all script (version 6.0) to identify pial and white matter surfaces. Processed anatomical data for each participant was used as the alignment target for all functional datasets which were kept within each participant’s native space. We used AFNI’s afni_proc.py to preprocess functional images, including motion correction (6-parameter affine transform), unwarping (using the forward/reverse phase-encode spin echo images), and coregistration (using the unwarped spin-echo images to compute alignment parameters to the anatomic target images).

We projected data to the cortical surface, then back into volume space, which incurs a modest amount of smoothing perpendicular to the cortical surface. To optimize distortion correction, we divided functional sessions into 3-5 subsessions, which consisted of 1-4 fMRI runs and a pair of forward/reverse phase encode direction spin echo images each, which were used to compute that subsession’s distortion correction field. For the spatial attention and mapping tasks, we did not perform any spatial smoothing beyond the smoothing introduced by resampling during coregistration and motion correction. For retinotopic mapping scans, we smoothed data by 5 mm FWHM on the surface before projecting back into native volume space.

### Region of interest definition

Using a previously described task that utilizes the voxel receptive field (vRF) method (Dumoulin & Wandell, 2008; Mackey et al., 2017; Thayer & Sprague, 2023), we identified 15 retinotopic ROIs for each participant (**Fig. 3A** shows example participant). Each run of the retinotopy task required participants to attend several random dot kinematograms (RDKs) within bars that would sweep across the visual field. Three equally sized segments were presented on each step, and the participants had to determine which of the two peripheral segments (opposite motion directions, always parallel to the long axis of the bar) matched the motion direction in the central segment with a button press. Participants received feedback via a red or green color change at fixation. We used a three-down/one-up staircase to maintain ∼80% accuracy throughout each run so that participants would continue to attend the RDK bars. RDK bars swept 17.5° of the visual field. Bar width and sweep direction were pseudo-randomly selected from several different widths (ranging from 2.0° to 7.5°) and four directions (left-to-right, right-to-left, bottom-to-top, and top-to-bottom).

We fit a vRF model for each voxel in the cortical surface (in volume space) using averaged and spatially smoothed (on the cortical surface; 5 mm FWHM) time series data across all retinotopy runs (7-12 per participant, 9.7 runs each on average). We used a compressive spatial summation isotropic Gaussian model (Kay et al., 2013; Mackey et al., 2017a) as implemented in a customized, GPU-optimized version of mrVista (see Mackey et al., 2017 for detailed description of the model). High-resolution stimulus masks were created (270 x 270 pixels) to ensure similar predicted responses within each bar size across all visual field positions. Model fitting began with an initial high-density grid search, followed by subsequent nonlinear optimization (Nelder-Mead simplex algorithm; bounded gradient descent within a specified range of best-fit parameters from grid search).

We visualized retinotopic maps by projecting vRF best-fit polar angle and eccentricity parameters with variance explained ≥10% onto each participant’s inflated cortical surfaces via AFNI and SUMA (**Fig. 3A**). We drew retinotopic ROIs (V1, V2, V3, V3AB, hV4, LO1, LO2, VO1, VO2, TO1, TO2, IPS0-3) on each hemisphere’s cortical surface based on previously-established polar angle reversal and foveal representation criteria (Amano et al., 2009; Mackey et al., 2017; Swisher et al., 2007; Wandell et al., 2007; Winawer & Witthoft, 2015). Finally, ROIs were projected back into volume space to select voxels for analysis. For analysis, we focused on groups of ROIs which share foveal confluences (as in Hallenbeck et al., 2021). Data from individual retinotopic regions were consistent with these aggregate ROIs, and are presented in **Figs. S3-4.**

### Inverted encoding model

To reconstruct neural priority maps based on each trial’s measured activation pattern within each ROI, we implemented an inverted encoding model (IEM) for spatial position. This analysis involves first estimating an encoding model [sensitivity profile over the relevant feature dimension(s) as parameterized by a small number of modeled information channels] for each voxel in a region by using a “training set” of data reserved for this purpose (spatial mapping runs). The encoding models across all voxels within a region are then inverted to estimate a mapping used to transform novel activation patterns from a “test set” (selective attention task runs) into activation in the modeled set of information channels. Adopting analysis procedures from previous work, we built an encoding model for spatial position based on a linear combination of spatial filters (Sprague et al., 2014, 2015; Sprague & Serences, 2013). Each voxel’s response was modeled as a weighted sum of 37 identically shaped spatial filters arrayed in a triangular grid (**Fig. 3B**). Centers were spaced by 2.83°, and each filter was a Gaussian-like function with full-width half-maximum of 3.12°:

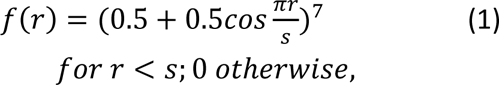

where *r* is the distance from the filter center and *s* is a “size constant” reflecting the distance from the center of each spatial filter at which the filter returns to 0. Values greater than this are set to 0, resulting in a single smooth round filter at each position along the triangular grid (*s* = 7.86°; see **Fig. 3B** for illustration of filter layout and shape; see also Sprague et al., 2014, 2015; Sprague & Serences, 2013).

This triangular grid of filters forms the set of information channels for our analysis. Each mapping task stimulus is converted from a contrast mask (1 for each pixel subtended by the stimulus, 0 elsewhere) to a set of filter activation levels by taking the dot product of the vectorized stimulus mask and the sensitivity profile of each filter. Once all filter activation levels are estimated, we normalize so that the maximum filter activation is 1.

Following previous reports (Brouwer and Heeger, 2009; Sprague and Serences, 2013a), we model the response in each voxel as a weighted sum of filter responses (which can loosely be considered as hypothetical discrete neural populations, each with spatial receptive fields (RFs) centered at the corresponding filter position):

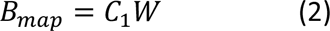

where B_map_ (*n* trials x *m* voxels) is the observed BOLD activation level of each voxel during the spatial mapping task (averaged over four TRs, 4.5 – 7.5 s after mapping stimulus onset), *C*_1_ (*n* trials x *k* channels) is the modeled response of each spatial filter, or information channel, on each non-target trial of the mapping task (normalized from 0 to 1), and *W* is a weight matrix (*k* channels x *m* voxels) quantifying the contribution of each information channel to each voxel.

Because we have more stimulus positions than modeled information channels, we can solve for *W* using ordinary least-squares linear regression:

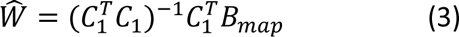

This step is univariate and can be computed for each voxel in a region independently. Next, we used all estimated voxel encoding models within a ROI (*Ŵ*) and a separate pattern of activation from the spatial attention task (each TR from each trial, in turn) to compute an estimate of the activation of each channel (*Ĉ*_2_; *n* trials x *k* channels), which gave rise to that observed activation pattern across all voxels within that ROI (*B*_*task*_; *n* trials x *m* voxels):

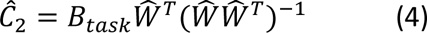

Once channel activation patterns are computed (*Eq. 4*), we compute spatial reconstructions by weighting each filter’s spatial profile by the corresponding channel’s reconstructed activation level and summing all weighted filters together. This step aids in visualization, quantification, and coregistration of trials across stimulus positions but does not confer additional information.

Because stimulus positions were unique on each trial of the selective attention task (**Fig. 2B; S1**), direct comparison of image reconstructions on each trial is not possible without coregistration of reconstructions so that stimuli appeared at common positions across trials. To accomplish this, we adjusted the center position of the spatial filters on each trial such that we could rotate the resulting reconstruction. For **Fig. 3C**, we rotated each trial such that one target (‘Stimulus 1’) was centered at x=4.22° and y=0° and the other stimulus (‘Stimulus 2’) was in the upper visual hemifield, which required flipping half of the reconstructions across the horizontal meridian. On Attend 1 trials, Stimulus 1 is defined as the cued (attended) stimulus and Stimulus 2 as the uncued (unattended) stimulus. On Attend 2 trials, both stimuli are cued (attended); Stimulus 1 is defined as the stimulus whose coherent spiral target appeared first (requiring a behavioral response), and Stimulus 2 as the stimulus whose target appeared second. On Attend fixation trials, neither stimulus is cued and target onset order is randomized across stimulus slots; the Stimulus 1/2 assignment follows the same positional convention used in the other conditions.

### Quantifying map activation

To quantify the strength of activation at stimulus location(s) in reconstructed neural priority maps, we averaged the pixels within each reconstruction located within a 1.27° radius disk centered at each stimulus’s known position. This gives us a single value for each stimulus location on each trial. We then sorted these measurements, which reflect linear transformations of BOLD activation levels, based on the contrast of the stimuli and the attention condition (**Figs. 3D and 4B**).

### Statistical analysis

For all statistical tests, we used parametric tests (repeated-measures ANOVAs and *t*-tests, where appropriate).

When analyzing BOLD responses, we first compared reconstructed map activation across stimulus conditions by computing separate 2-way repeated-measures ANOVAs for each ROI, with relevance condition (Attend fixation, Attend 1, Attend 2) and matched stimulus contrast (20%, 40%, 80%) as factors (**Fig. 3D; Table S1**). To test whether relevance-related activation in neural priority maps depends on the number of relevant location(s), we first visualized the difference between map activation profiles between each stimulus-relevant location (Attend 1, Attend 2) and the Attend fixation condition at each contrast (**Fig. 4A** shows an example contrast). Then, we quantified this difference score at the location of Stimulus 1, which was relevant in each condition, at each contrast (**Fig. 4B**). We performed a 2-way repeated-measures ANOVA on each ROI with factors of relevance condition (Attend 1, Attend 2) and matched stimulus contrast (20%, 40%, 80%). Additionally, we conducted pairwise t-tests within each ROI at each matched contrast between relevance conditions (Attend 1 vs Attend 2). Results presented in the manuscript have not been corrected for multiple comparisons. However, note that the primary conclusions are an inability to reject the null hypothesis; correction is not performed to avoid inflating type II error rates.

### Glitches

Due to an indexing error in the stimulus presentation script, only a single target event was presented on trials where Stimulus 1’s contrast was 40% or 80%: on these trials, Stimulus 2 never contained a coherent spiral target, regardless of attention condition or Stimulus 2’s contrast. Because this occurred equally frequently across all task conditions, and because the target location on Attend 2 trials was still fully unpredictable (Stimulus 1 always contained the only target, but its spatial location was randomized trial-to-trial), this bug is not expected to impact the interpretation of the results. Additionally, behavioral response (button press) data was not saved for a small subset of trials for a subset of participants (maximum of 12 missing trials total per participant). These trials are not included in behavioral data analyses.

### Data & code availability

[Upon publication], all data & code necessary to reproduce results of this study [will be made] available at [URL ADDED AFTER ACCEPTANCE] and https://github.com/SpragueLab/Pri2StimDist/.

## Results

We measured blood oxygenation level-dependent (BOLD) activation patterns from several independently identified retinotopic regions using fMRI while participants covertly attended to one (Attend 1) of two visual stimuli (two patches of randomly oriented dark and light lines both presented at 20%, 40%, or 80% contrast), to both stimuli simultaneously (Attend 2), or to the fixation point (Attend fix; **Fig. 2A-B**). Participants identified the category of a brief target stimulus (coherent lines that formed a spiral) at the cued location in the Attend 1 condition, or in the case of the Attend 2 condition, identified the category of the target stimulus that appeared first. In the Attend fixation condition, participants identified whether the horizontal or vertical arm of a flickering fixation cross lengthened.

Behavioral performance inside the scanner was consistent with a modest cost to monitoring multiple stimuli simultaneously (**Figs. 2C-E**). Based on a two-factor repeated-measures ANOVA (factor 1: relevance condition, Attend 1 and Attend 2; factor 2: matched stimulus contrast: 20%, 40%, 80%), response accuracy significantly differed as a function of task demands (F(1,9) = 7.048, *p* = 0.026, partial η² = 0.439) and stimulus contrast (F(2,18) = 6.445, *p* = 0.008, partial η² = 0.417), but not their interaction (F(2,18) = 0.574, *p* = 0.573, partial η² = 0.060). Follow-up paired t-tests for each stimulus contrast show a significant difference between attend-one and attend-two trials for 20% contrast stimuli (t(9) = 3.10, *p* = 0.013), but no significant difference for the other contrasts (40%: t(9) = 0.76, *p* = 0.466; 80%: t(9) = 1.74, *p* = 0.116). For RT, the same ANOVA supported no significant main effect of task demands (F(1,9) = 1.366, *p* = 0.272, partial η² = 0.132), stimulus contrast (F(2,18) = 3.248, *p* = 0.062, partial η² = 0.265), nor a significant interaction (F(2,18) = 1.148, *p* = 0.340, partial η² = 0.113). Pairwise t-tests for each contrast found no significant differences between attention conditions (all *p* ≥ 0.100). Finally, for average target coherence (which was manually adjusted for each contrast and task condition between runs with the goal of ∼80% accuracy), there were no significant effects of task demands (F(1,9) = 1.528, *p* = 0.248, partial η² = 0.145), stimulus contrast (F(2,18) = 2.676, *p* = 0.096, partial η² = 0.229), nor an interaction (F(2,18) = 2.067, *p* = 0.156, partial η² = 0.187). Pairwise t-tests for each contrast found no significant differences between attention conditions (all *p* ≥ 0.147). Slightly lower accuracy on trials with multiple attended locations (**Fig. 2C**) is consistent with a modest behavioral cost to monitoring multiple locations simultaneously, aligning with aspects of previous behavioral and neuroimaging work (e.g., Harrison et al., 2023; McMains & Somers, 2005; Pestilli et al., 2011). Altogether, our in-scanner behavioral results demonstrate slightly less accurate performance when multiple locations were relevant for behavior, motivating our efforts to characterize neural mechanisms for selecting multiple relevant objects.

Next, we sought to determine how neural priority maps implemented as activation profiles across retinotopic regions of cortex index task relevance of stimulus locations. If task-relevance of a covertly-attended location drives neural response modulations among neural populations tuned to the relevant location(s), then we expect to see relevance-related activation at the relevant location(s) on the priority map, even when stimuli are identical across relevance conditions If there are neural constraints limiting the simultaneous and equivalent signaling of multiple locations as relevant, activation at a stimulus location will be strongest when only a single stimulus is task-relevant, with weaker activation when multiple stimuli are made relevant. Instead, if relevance-related activation primarily serves to enhance the propagation of information out of relevant cortex for further processing but is not itself limited, we expect to see similar relevance-related map activation at stimulus locations across our task relevance conditions.

To visualize and quantify neural priority maps encoded by activation profiles over retinotopically-organized regions, we used a multivariate fMRI image reconstruction technique (an inverted encoding model, IEM) (Sprague & Serences, 2013). This method projects activation patterns measured in voxel space within a brain region to a modeled spatial coordinate system (retinotopic position), and has been used in previous studies characterizing the impact of stimulus and task features on neural priority map profiles (Sprague et al., 2018; Sprague & Serences, 2013; Thayer & Sprague, 2023, 2025). First, we estimated the spatial sensitivity profile of each voxel within our retinotopically-defined regions of interest (ROIs; **Fig. 3A**) using data measured from a separate set of “mapping” scans. We then used the resulting sensitivity profiles (which are discretized versions of ‘population receptive fields’; Dumoulin & Knapen, 2018; Dumoulin & Wandell, 2008; Wandell & Winawer, 2015) across all voxels to reconstruct a map of retinotopic space in visual field coordinates from single-trial activation patterns measured during the main task (**Fig. 3B**). From these maps, we quantified activation strength at each stimulus location to evaluate how neural priority maps index the relevance of multiple stimulus locations.

First, we compared neural priority maps estimated from visual field map clusters in visual, dorsal, lateral, and ventral regions (**Fig. 3C**). For illustration, we focus on trials where stimuli appeared 144° apart (see **Fig. 2B**). Qualitatively, activation is visible at multiple stimulus locations in most ROIs, especially in the Attend 2 condition. Most ROIs show a scaling in map activation at stimulus locations with an increase in stimulus contrast, consistent with sensitivity to stimulus salience regardless of task demands (and replicating Sprague et al, 2018), with the exception of LO1/LO2. On Attend 1 trials, where both stimuli are identical but only Stimulus 1 is task relevant, all ROIs show a clear augmentation of map activation at the task-relevant location as compared to either the same location when irrelevant (Attend fixation), or the other task-irrelevant stimulus location on the same trials. Critically, on Attend 2 trials, where both stimuli are now task-relevant, map activation is stronger at both locations as compared to Attend fixation, and is not visibly different from map activation on Attend 1 trials. That is, the modulatory impact of task relevance appears binary: when a stimulus is cued to be task relevant, its location is more active in neural priority maps across the visual system.

To quantify these effects, we extracted the mean activation in each reconstructed map at the position of Stimulus 1 from each task relevance and stimulus contrast condition and plotted contrast response functions for each ROI and condition (**Fig. 3D**). To determine whether map activation scaled with stimulus contrast, task relevance condition, or their interaction, we performed repeated-measures ANOVAs for each ROI with factors of task relevance condition (Attend fixation, Attend 1, Attend 2) and matched stimulus contrast (20%, 40%, 80%). We found significant main effects of task relevance condition across all ROI clusters (*p* ≤ 0.038, maximum *p*-value TO1/TO2), and significant main effects of stimulus contrast in nearly all ROI clusters (*p* ≤ 0.026, maximum p-value TO1/TO2), with LO1/LO2 the lone cluster with no significant effect (p = 0.653). There was no interaction between condition and contrast in any visual area that we evaluated (*p* ≥ 0.455, minimum *p*-value in LO1/LO2; all statistics for 2-way ANOVA for each ROI available in **Table S1;** Results from Stimulus 2 are shown in **Fig. S2**; Results from all individual retinotopic ROIs shown in **Fig. S3**). As illustrated by the contrast response functions plotted in **Fig. 3D**, the main effect of task relevance condition is primarily driven by a difference between trials where peripheral stimuli are irrelevant (Attend fix) and trials where the peripheral stimulus is relevant (Attend 1 and Attend 2), with minimal difference between the contrast response functions for Attend 1 and Attend 2 trials. (note that activation for Stimulus 1 is plotted, which is the task-relevant stimulus on Attend 1 trials. See **Fig. S2** for results for Stimulus 2). Additionally, the lack of observed interaction between task relevance and contrast in any ROI is consistent with previous fMRI studies establishing a primarily additive effect of covert visual attention on neural responses (Buracas and Boynton, 2007; Murray, 2008; Pestilli et al., 2011; Itthipuripat et al., 2019a).

Based on the substantial effect of task relevance on activation levels in neural priority maps at stimulus locations, we next sought to determine if the activation difference due to task relevance was equivalent between the Attend 1 and Attend 2 conditions. That is, does cueing multiple locations as equally task-relevant result in a different strength of activation at neural priority maps than when a single location is cued? First, we visualized the selective impact of task relevance on neural priority maps from each ROI cluster by computing the difference map between Attend 1 – Attend fix and Attend 2 – Attend fix (**Fig. 4A**), which we interpret as a ‘relevance map’ for Attend 1 and Attend 2 conditions, respectively. Qualitatively, several features stand out: first, the impact of task relevance is relatively focal at the task-relevant location(s), with minimal spread over the intervening area of the visual field. This is consistent with previous observations that humans and primates can effectively attend to disjoint locations (McMains and Somers, 2004, 2005; Niebergall et al., 2011). Second, the relevance signal (red blob) is approximately equally strong at the location of Stimulus 1 in each condition’s difference map, consistent with these maps encoding each location’s relevance state in a binary fashion. Finally, in several ROI clusters, especially IPS2/IPS3 and LO1/LO2, cueing a peripheral location as task-relevant (and the fixation point as task-irrelevant) results in a shifting of the relevance profile, with evidence for a qualitative reduction of activation near the fixation point corresponding to an increase in activation at the relevant stimulus location(s) (this is also visually apparent in the condition-wise neural priority maps shown in **Fig. 3C**).

To quantify whether neural priority maps index relevant location(s) at equivalent levels across task relevance conditions (Attend 1, Attend 2), we computed an index of relevance-related map modulation for each condition by subtracting activation in the Attend fixation condition (where neither stimulus was relevant) from that in the Attend 1 and Attend 2 conditions for each contrast level (**Fig. 4B**). This allowed us to directly compare the strength of relevance-related activation change across conditions. If the degree of activation associated with a relevant stimulus location is independent of the relevance of other stimulus locations, then we would expect to find no significant difference in modulation of map activation at the Stimulus 1 location between Attend 1 and Attend 2 conditions. Indeed, we observed no consistent evidence of an effect of task relevance condition on relevance-related changes in map activation. Results from a repeated-measures 2-way ANOVA (factors of task relevance condition: Attend 1, Attend 2, and matched stimulus contrast: 20%, 40%, 80%) showed that most ROI clusters had no main effect of relevance condition (p ≥ 0.072, minimum non-significant p-value V3AB), with the exception of midlevel extrastriate clusters hV4/VO1/VO2 (p = 0.034) and LO1/LO2 (*p* = 0.002). Follow-up paired t-tests between activation differences from Attend 1 and Attend 2 at each contrast within these ROI clusters failed to identify consistent modulations across contrasts: hV4/VO1/VO2 did not have any significant differences for individual contrasts (*p* ≥ 0.182), and activation difference in LO1/LO2 only significantly differed at 40% contrast (*p* = 0.037). Activation differences did not vary as a function of contrast within any ROI clusters (*p* ≥ 0.355, minimum *p*-value TO1/TO2), and there were no significant interactions between task relevance condition and contrast (*p* ≥ 0.214, minimum *p*-value V1/V2/V3). All statistics are available in **Table S3**, and results from individual retinotopic ROIs (which are consistent with those reported here) are shown in **Fig. S4**. Altogether, these results suggest that activation profiles in neural priority maps measured across a substantial portion of the human visual system index whether an individual location is task-relevant, with no impact of the relevance of other locations.

## Discussion

We evaluated how neural priority maps in retinotopic cortex (**Fig. 1**) encode the relevance of stimulus locations while participants performed a demanding task requiring covert spatial attention to one or two stimuli. While behavioral performance showed only a modest cost of dividing attention across task conditions (**Fig. 2**), neural priority maps reconstructed from fMRI activation profiles showed clear sensitivity to whether a stimulus location was relevant, independent of whether another location was also relevant (**Figs. 3-4**). Moreover, the spatial profile of task-related modulation in neural priority maps was disjoint and localized to stimulus locations rather than conjoined across the relevant positions (**Fig. 4**). We interpret this pattern as reflecting a spatial “relevance map” that can independently select multiple stimulus locations to implement attentional modulations in neural activity, including changes in firing rates, correlation structure, or inter-regional communication. These data suggest that relevance-related modulations remain largely unconstrained when two locations must be simultaneously monitored. We propose that such modulations serve to enable further processing of relevant stimuli by downstream regions, where performance limitations may instead arise.

**Figure 1:**
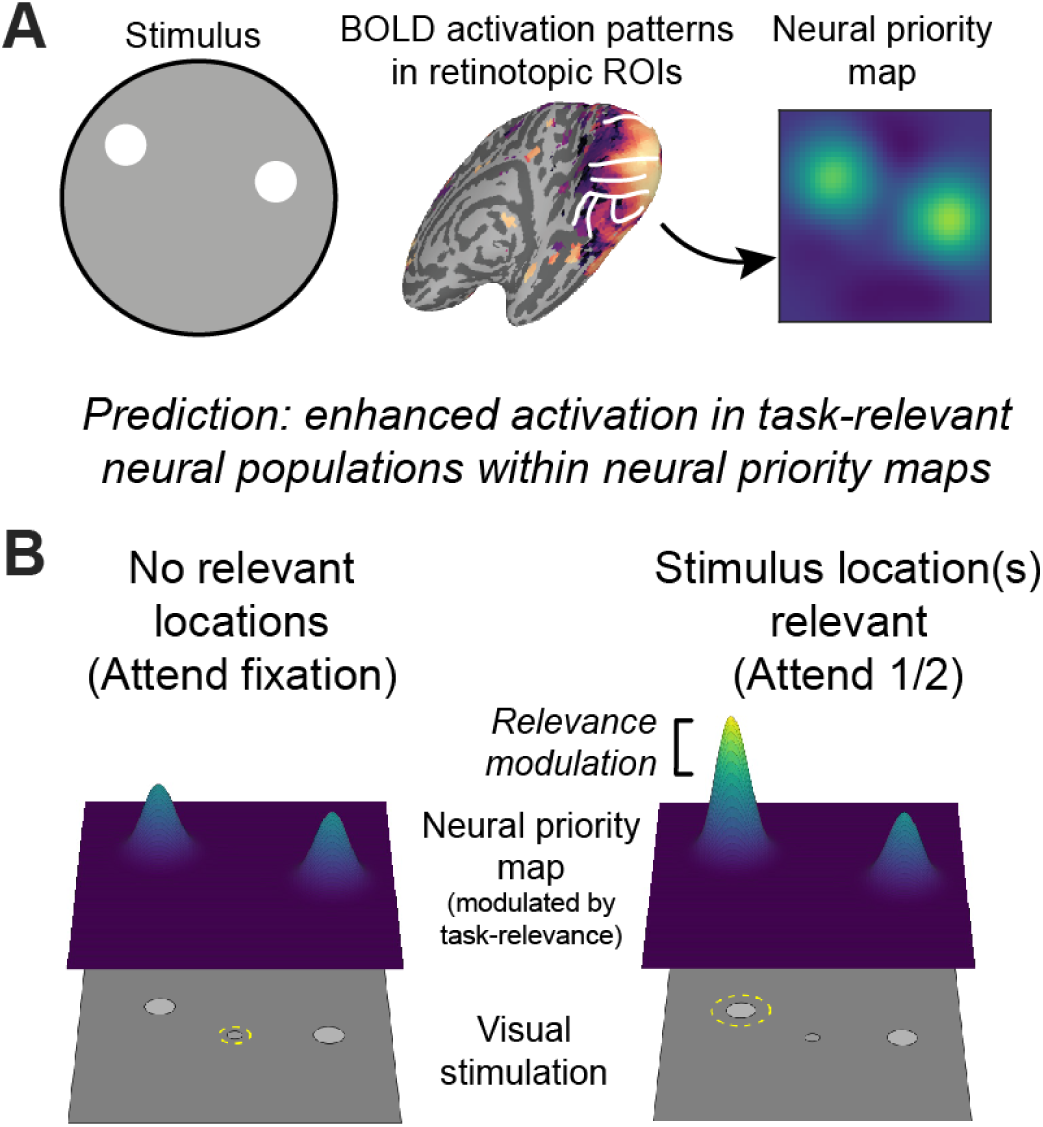
Identifying relevance maps in human cortex. **(A)** Neural priority maps are maps of the visual field with activity scaling according to the stimulus-based salience and/or task-based relevance of each location. Previous studies have reconstructed neural priority maps from activation patterns measured in retinotopic cortex (e.g., Sprague et al., 2018; Sprague & Serences, 2013) (see Fig. 3). **(B)** A relevance map is revealed as the modulatory profile of neural priority maps when task relevance only is manipulated (e.g., comparing trials where 1 stimulus is attended to trials where the fixation point is attended). Cartoon depiction of predicted result for Attend 1 condition shown. Yellow dashed lines in stimulus diagrams indicate attended location.

**Figure 2.**
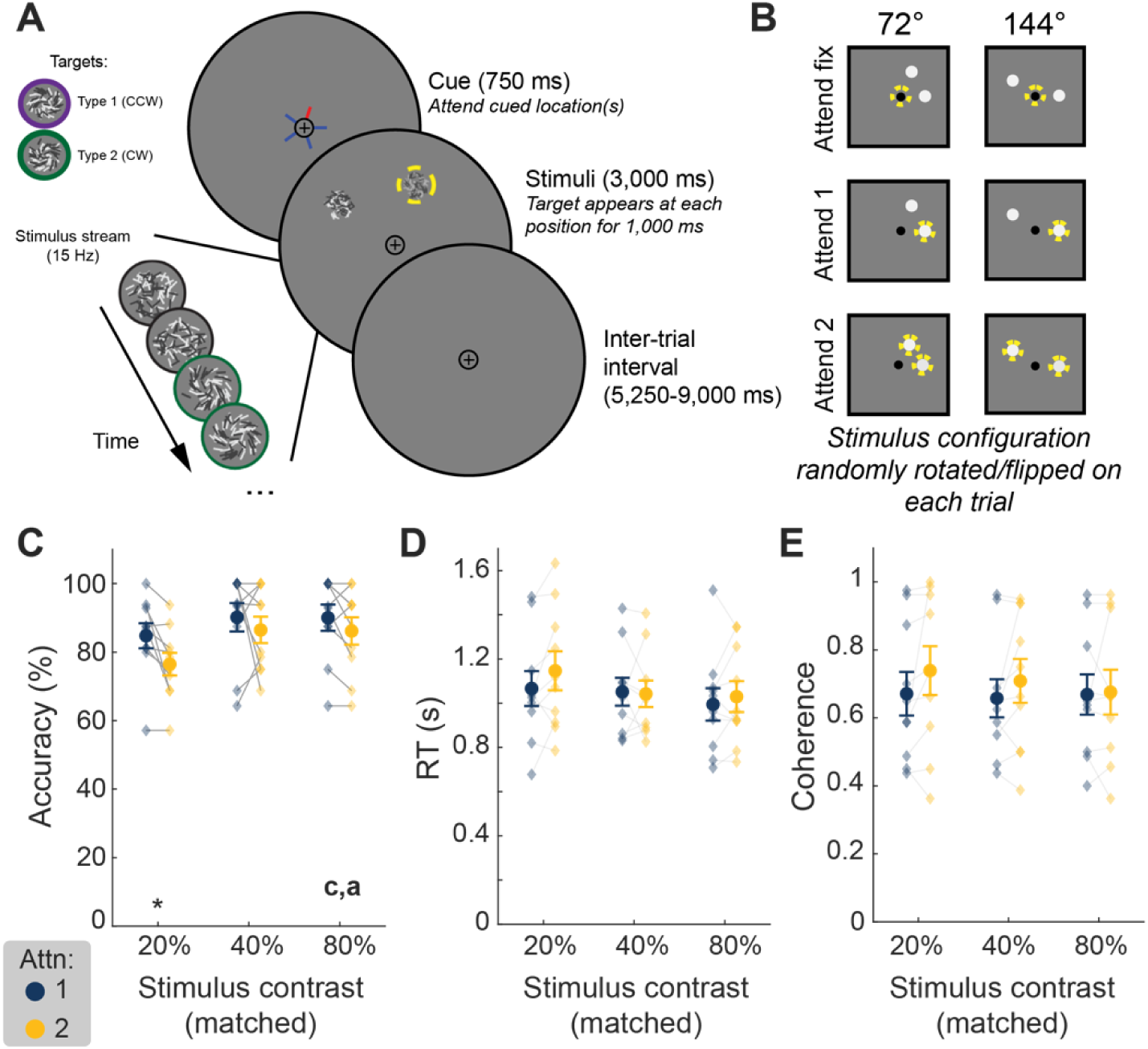
Task design and behavioral performance during fMRI scanning. **(A)** On each trial, participants (n = 10) saw a cue indicating with 100% validity whether the task-relevant stimulus was the fixation point (‘attend fix’), one peripheral stimulus (indicated by red arm), or both peripheral stimuli (two red arms). Participants monitored the cued stimulus/stimuli for a target event (Attend fix: increase in height/width of fixation cross; Attend 1 or 2: appearance of coherent structure in random line stimulus) and used a button box to perform a discrimination judgment of the target type (Attend fix: tall/wide fixation; Attend 1: CCW/CW target in single cued stimulus, Attend 2: CCW/CW target that appears first among the two stimuli). Example trial from Attend 1 condition shown; relevant stimulus highlighted by yellow dashed line (not shown to participants) **(B)** Stimuli appeared at a fixed eccentricity and random rotation around fixation, drawn from two arrangements. Both stimuli appeared at the same contrast on each trial to isolate changes in neural activation profiles associated with task-relevance. See **Figure S1** for additional design details, including trial conditions not reported in the present study. **(C)** Overall task accuracy when peripheral stimulus/stimuli were relevant for each stimulus contrast. **(D)** Response time (as in C). **(E)** Mean coherence across fMRI sessions. Coherence was adjusted for each condition across scanning runs to attain ∼80% performance; higher coherence results in an easier task. For each of **Fig. 2C-E**, we performed a 2-way repeated-measures ANOVA with factors of relevance condition (Attend 1, Attend 2) and stimulus contrast (20%, 40%, 80%). ‘a’ indicates significant main effect of relevance condition; ‘c’ indicates significant main effect of contrast; ‘x’ indicates significant interaction. ‘*’ indicates significant paired t-test between relevance conditions at each individual contrast for each analysis. Significance markers shown at *p* ≤ 0.05, uncorrected. Error bars standard error of the mean (SEM); individual data points shown as connected diamonds.

**Figure 3.**
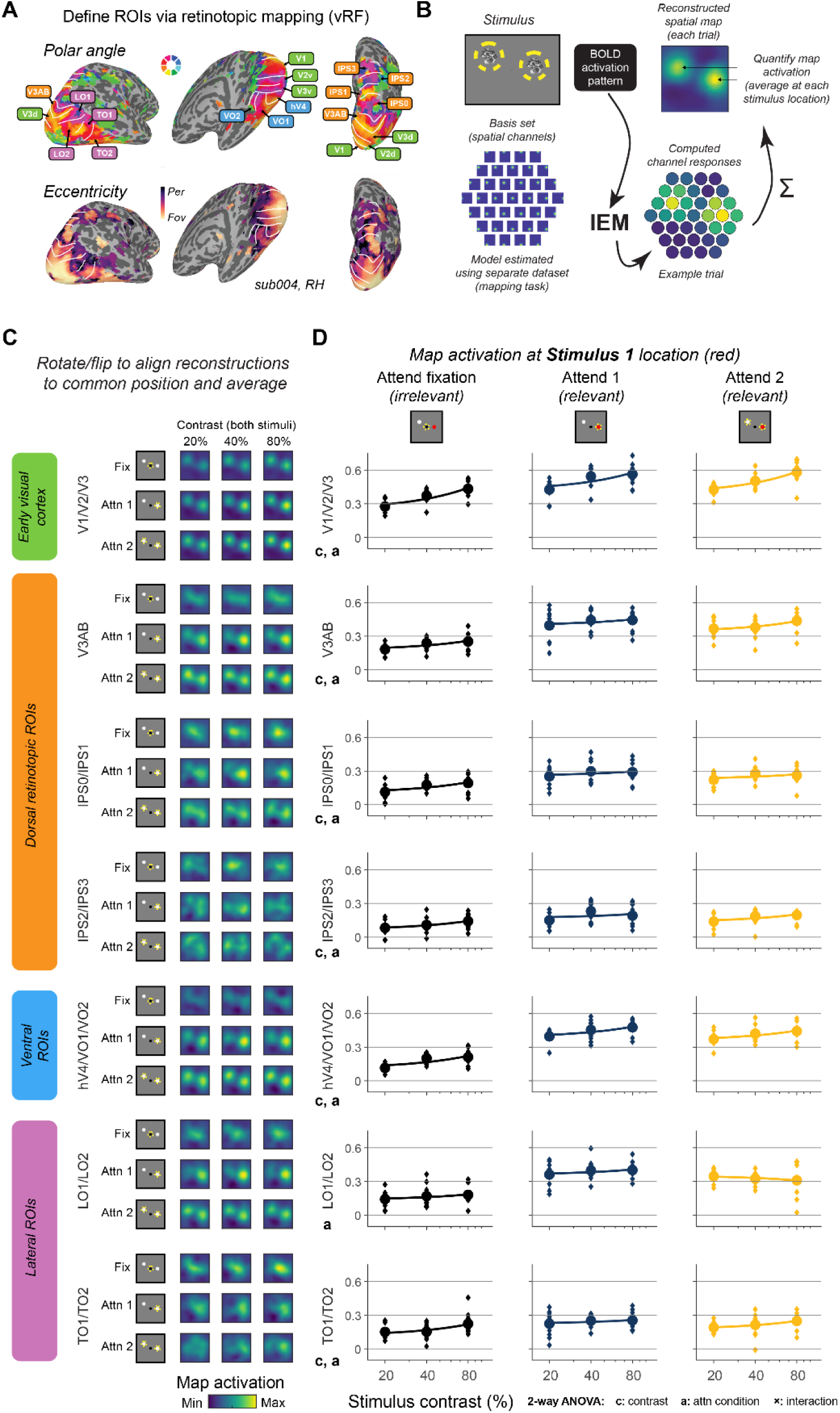
Neural priority maps jointly index multiple relevant locations. **(A)** We used voxel receptive field (vRF)-based retinotopic mapping techniques to identify visual field map clusters in each participant’s visual, parietal, and temporal cortex. Example participant’s polar angle and eccentricity maps presented (voxelwise threshold: ≥ 10%) on their inflated right hemisphere with ROI borders highlighted. **(B)** We reconstructed neural priority maps (activation profiles in screen coordinates) from measured fMRI activation patterns across each joint ROI using an inverted encoding model (Sprague and Serences, 2013a; Sprague et al., 2018a). This method estimates an encoding model for each voxel using a separate dataset (not shown), then inverts this encoding model to transform a new measured activation pattern to activation profiles across modeled information channels, which can be aligned across similar trials and visualized as neural priority maps. See *Methods*. **(C)** Reconstructed neural priority maps for each condition on 144 deg stimulus separation trials (stimulus cartoon denoting relevant location(s) with yellow circles on left). Qualitatively, both stimuli are readily apparent in most maps (especially at higher contrast and when both stimuli are relevant). Colormap is scaled independently across all maps within each ROI. **(D)** Quantified map activation at location of Stimulus 1, which is always a relevant location when a peripheral stimulus is cued (see cartoon). Activation is quantified as mean map activation at map pixels corresponding to stimulus location on each trial. See **Figure S2** for similar quantification of Stimulus 2, and **Figure S3** for similar quantification for each retinotopic ROI individually. When a stimulus is made relevant via a cue to attend that stimulus or both stimuli, its activation is equivalently enhanced across all contrast values. Results from 2-way repeated-measures ANOVA with factors of task condition (attend fix, attend 1, attend 2) and matched stimulus contrast (20%, 40%, 80%) indicated with ‘c’ for main effect of contrast, ‘a’ for main effect of task condition, and ‘x’ for interaction at uncorrected p ≤ 0.05. See **Table S1** for all statistics. Circles and error bars are mean and SEM across n = 10; diamonds are individual participant datapoints. Line is linear function fit to mean (plotted log scale).

Most human studies examining task relevance compare neural responses to an attended stimulus with responses to unattended stimuli or to identical displays in which the stimulus is task-irrelevant (e.g., passive viewing or fixation tasks). These studies consistently show that attended stimuli evoke stronger neural responses, including increased BOLD activation (Gandhi et al., 1999; Kastner et al., 1999; Buracas and Boynton, 2007; Murray, 2008; Pestilli et al., 2011), enhanced representations in reconstructed spatial priority maps (Sprague and Serences, 2013a; Sprague et al., 2018a; Thayer and Sprague, 2025), and changes in voxel receptive field encoding properties (Çukur et al., 2013; Klein et al., 2014; Kay et al., 2015; Sheremata and Silver, 2015; Vo et al., 2017; van Es et al., 2018; You et al., 2025). Our results are consistent with this literature: activation in reconstructed priority maps was stronger at relevant than irrelevant stimulus locations in the Attend 1 condition, and when peripheral stimuli were task-relevant compared to trials in which attention was directed to fixation (**Fig. 3**).

While studies of attention to a single stimulus produce broadly consistent results, studies requiring attention to multiple stimuli have yielded more variable findings. Some fMRI studies report reduced attentional modulation when attention is divided across multiple stimuli compared with when a single stimulus is attended (McMains and Somers, 2005; Pestilli et al., 2011; Tünçok et al., 2025), while others report little or no reduction in attentional modulation under distributed attention (White et al., 2017, 2019; Zhang et al., 2017; Narhi-Martinez et al., 2025). fMRI studies which aim to directly measure the visual field or cortical extent of attentional modulation provide further support that retinotopic regions can encode the set of spatial locations relevant for a task (Bloem et al., 2025; Herrmann et al., 2010; Itthipuripat et al., 2014, 2019; Park & Serences, 2022; Puckett & DeYoe, 2015; Tünçok et al., 2025). Electrophysiological studies in nonhuman primates show similarly mixed results: some show evidence that cue validity scales spike rates in extrastriate neurons (Mayo and Maunsell, 2016), while others see similar spike rates regardless of the number of attended stimuli in several visual regions (Arcizet et al., 2018; Denfield et al., 2018). Notably, a study employing voltage-sensitive dye measurements in macaque primary visual cortex – a technique sensitive to a combination of local activity and input to a region – during a cued attention task revealed equivalent enhancement at cued stimulus locations whether one or four locations were behaviorally relevant (Chen and Seidemann, 2012). Our results align with these latter findings, showing that attending to multiple locations produces levels of attentional enhancement comparable to those observed when a single location is attended (**Figs. 3-4**).

**Figure 4.**
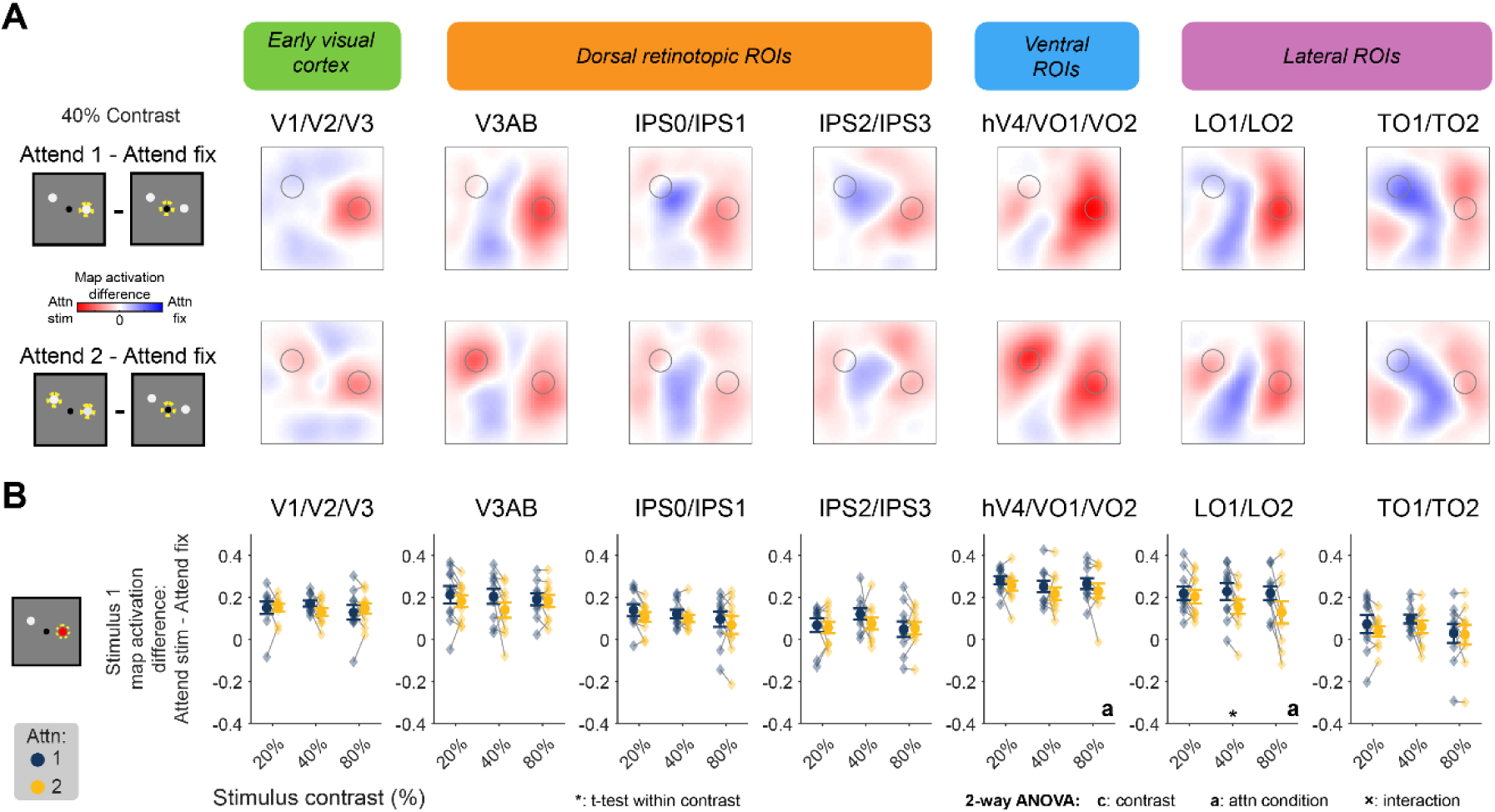
Attention-related modulations in neural priority maps reveal a relevance map. **(A)** Difference maps computed between conditions where peripheral location(s) were relevant (Attend 1; Attend 2) and when they were irrelevant (Attend fix; see cartoons). Red and blue areas indicate locations where task-relevance increases/decreases map activation, respectively. Gray circles indicate stimulus locations. Maps averaged across all participants (n = 10) for trials from an example contrast (40%). Color scale is matched across all maps and centered at zero. **(B)** Relevance-related map modulation quantified for each relevance condition, contrast, and ROI. Circles/error bars plot mean/SEM and connected diamonds plot individual participant data. No regions show consistent differences between strength of relevance-related map modulation across task conditions across all contrasts. Results from 2-way repeated-measures ANOVA with factors of relevance condition (attend 1, attend 2), matched stimulus contrast (20%, 40%, 80%), and their interaction are shown for significant comparisons at *p* ≤ 0.05 (uncorrected). Significant results from pairwise t-tests between relevance conditions at each contrast are shown with ‘*’ (*p* ≤ 0.05, uncorrected). Full statistical results are available in **Table S3.**

### Myriad neural mechanisms of covert spatial attention

A growing body of work in model species has identified several mechanisms by which covert spatial attention can influence sensory processing. In addition to classical changes in neural gain (e.g., Luck et al., 1997), attention has been linked to changes in synaptic efficacy (Briggs et al., 2013; Hembrook-Short et al., 2019), correlation structure (Cohen and Maunsell, 2009), and inter-areal communication (Fries et al., 2001). Importantly, many of these mechanisms do not necessarily predict that the *magnitude* of attentional modulation should scale with the number of task-relevant stimuli. That is, once a stimulus location is designated as behaviorally relevant, neural populations representing that location may be selected for prioritized processing regardless of how many other locations are concurrently relevant. Under such an account, multiple stimulus locations could be selected to a similar degree, with limits on behavioral performance arising at later stages of processing or decision-making rather than through weakened early sensory representations (Bay and Wyble, 2014; White et al., 2017, 2019; Moreland et al., 2020; Harrison et al., 2023).

Even in cases where the magnitude of attentional modulation does change, many proposed neural mechanisms do not predict a corresponding monotonic change in overall BOLD activation. For example, changes in neural variability or neural synchrony that leave mean firing rates unchanged could yield similar BOLD responses despite measurable differences in electrophysiological activity. Consistent with this dissociation, a prior study directly comparing EEG and fMRI during covert spatial attention demonstrated that fMRI BOLD signals and asynchronous electrophysiological measures such as lateralized alpha activity primarily track spatial relevance, whereas evoked and synchronous electrical signals—such as ERP components or SSVEP power at the stimulus frequency—vary in their attentional modulation with stimulus contrast (Itthipuripat et al., 2019). These findings mirror intracranial observations revealing a complex and sometimes indirect relationship between local circuit dynamics and measured BOLD responses in visual cortex (Logothetis et al., 2001; Sirotin and Das, 2009; Winawer et al., 2013; Hermes et al., 2017).

### Relevance maps throughout retinotopic cortex

Our results suggest that the neural signatures of spatial attention observed here are more consistent with a relevance-based selection signal than with graded enhancement of stimulus-evoked responses limited by the number of relevant stimuli. While some established mechanisms of covert attention—such as response or contrast gain increases—may be expected to introduce inter-stimulus interference through normalization or interneuron-mediated inhibition (Reynolds and Heeger, 2009; Carandini and Heeger, 2011), other mechanisms, including changes in synaptic efficacy, correlation structure, inter-areal synchrony, or subspace alignment, need not result in changes to overall activity dependent on the number of relevant locations. As a result, these mechanisms could plausibly support equivalent attention-related BOLD modulations across one or multiple selected locations despite detectable differences in electrophysiological measures (Winawer et al., 2013; Hermes et al., 2017; Itthipuripat et al., 2019a). We argue that such BOLD signal modulations in retinotopic cortex reveal the neural signature of the ‘relevance map’ (**Fig. 1**), which highlights the known-to-be relevant stimulus location(s) among all possible locations in the display for further processing. These relevance-related modulations in retinotopic cortex are likely driven by input from higher-order frontal and parietal regions involved in sculpting signals in earlier regions (Moore et al., 2003; Moore, 2006; Baldauf and Desimone, 2014; Moore and Zirnsak, 2017; Li et al., 2025), consistent with an architecture in which top-down selection signals act independently on each task-relevant location.

### Comparisons to Visual WM Studies

Our observation of equivalent relevance-related BOLD signal modulation across multiple attended locations contrasts with previous demonstrations of load-related reductions in representations of multiple remembered locations in similar ‘relevance maps’ measured from fMRI activation patterns during visual spatial working memory (WM) studies (Sprague et al., 2014, 2016). In such studies, multiple spatial locations must be precisely maintained across a delay period to guide a later behavioral response while participants maintain fixation and view a blank screen. In this context, reconstructed neural priority maps in retinotopic regions across the dorsal stream show spatial WM representations which scale inversely with WM load (representations are weaker for two remembered locations than for one) and are related to behavioral performance. In the present study, we expected similar reductions in task-relevance-related signals as more locations were cued to be relevant. Instead, our finding that stimulus-evoked signals are equivalently modulated by task relevance suggests a complex interplay between top-down signals indicating that locations are task-relevant and bottom-up stimulus drive. Given that attention and WM are broadly thought to be tightly coupled processes sharing overlapping neural substrates (Awh and Jonides, 2001; Awh et al., 2006; Myers et al., 2015; Ede and Nobre, 2023), the divergence between our results and WM load studies is striking, and may hinge on the presence of ongoing visual input: when stimuli are present, activation modulations related to top-down relevance signals may be similar at each relevant stimulus location, whereas during blank delay periods, maintenance of multiple representations may reveal limitations in relevance-related activation absent simultaneous stimulus drive (Ekstrom et al., 2008). For example, tasks requiring a post-cued behavioral response entail some storage in visual WM (e.g., Tünçok et al., 2025), as do two-interval forced-choice contrast discrimination tasks (e.g., Pestilli et al., 2011). Future work manipulating task components—including visual WM demands, attended feature(s), stimulus intensity, and number of attended stimuli—will further illuminate when task-related modulations are constrained by load and when they are unconstrained, as observed here.

### Alternative explanations and limitations

Although these relevance-related BOLD modulations differ from predictions derived from psychophysical studies comparing valid and neutral cues during difficult visual discriminations, they may not directly reflect the quality of stimulus information encoded at relevant locations. We did not measure neural information about the discriminandum—the type of target stimulus—itself, differing from studies in which stimulus contrast was the task-relevant stimulus feature (Jehee et al., 2011). In these studies, attention-related activation modulations decreased at each of multiple simultaneously-attended locations compared to a single attended location (Pestilli et al., 2011; Hara et al., 2014; Itthipuripat et al., 2014a). Future studies jointly quantifying map-level activation profiles and trial-level information about task-relevant features (like orientation) would better dissociate modulations at the ‘source’ of top-down relevance signals from modulations of task-relevant information content in neural population codes (Li et al., 2025). More broadly, it remains possible that other task designs, stimulus configurations, or measurement modalities would reveal reductions in relevance-related enhancement as more locations become task-relevant. Factors including the number of attended locations, attended features (Chapman and Störmer, 2023), cue validity (Mayo and Maunsell, 2016), and requirements for storage in visual short-term or working memory (as in post-cue or two-interval forced-choice tasks; Luck et al., 1994; Palmieri & Carrasco, 2024; Pestilli et al., 2011; Pestilli & Carrasco, 2005; Tünçok et al., 2025) may all influence whether relevance-related modulations scale with number of relevant locations. Identifying these boundary conditions is an important direction for future work.

### Conclusions

Altogether, our results reveal that top-down modulation of neural priority maps operates independently at each task-relevant location, undiminished by the relevance of another simultaneously attended stimulus. We interpret this modulation profile as the neural manifestation of a ‘relevance map’, which we speculate reflects the populations within retinotopic cortex where the myriad neural mechanisms shown to support covert spatial attention are implemented. Per this framework, behavioral limits when attending to multiple items may arise from constraints in readout or decision processes rather than solely from reduced modulation of sensory representations. These findings open new questions about how the visual system encodes graded levels of relevance across locations, how electrophysiological signatures of this selection signal relate to the BOLD modulations observed here, and at what stage of processing the well-established behavioral costs of divided attention ultimately arise.

## Acknowledgments

This work was supported by Cooperative Agreement W911NF-19-2-0026 for the Institute for Collaborative Biotechnologies, a University of California, Santa Barbara Academic Senate Research Grant, NIH/NEI R01-EY035300, and an Alfred P Sloan Research Fellowship to T.C. Sprague. We thank Barry Giesbrecht, Miguel Eckstein, and Regina Lapate for helpful comments on the project at earlier stages.

## Supplemental material

**Figure S1.**
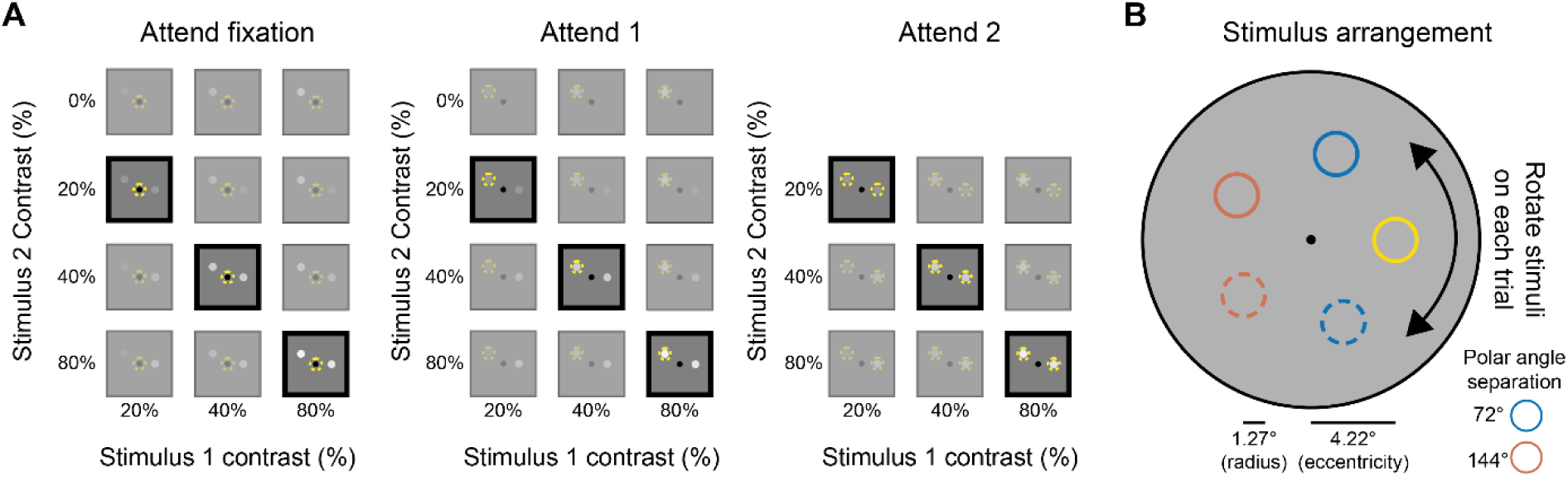
Full stimulus configuration and trial conditions. *(A)* Full set of trial conditions for each task relevance condition (Attend fixation, Attend 1, Attend 2). Each cell depicts the contrast pairing of Stimulus 1 (x-axis; 20%, 40%, 80%) and Stimulus 2 (y-axis; 0%, 20%, 40%, 80%) for a given trial type. Trials with Stimulus 2 contrast of 0% are single-stimulus trials in which only Stimulus 1 was presented. Cells outlined in black indicate matched-contrast two-stimulus trials; only these trials are included in the analyses reported in the present study. *(B)* Stimuli appeared at 4.22° eccentricity on each trial, drawn from two possible polar angle separations (72°, blue; 144°, red). Stimulus 1 served as the positional reference from which Stimulus 2’s location was derived. The entire stimulus arrangement was rotated randomly about fixation on each trial, such that each trial involved a unique stimulus configuration.

**Figure S2.**
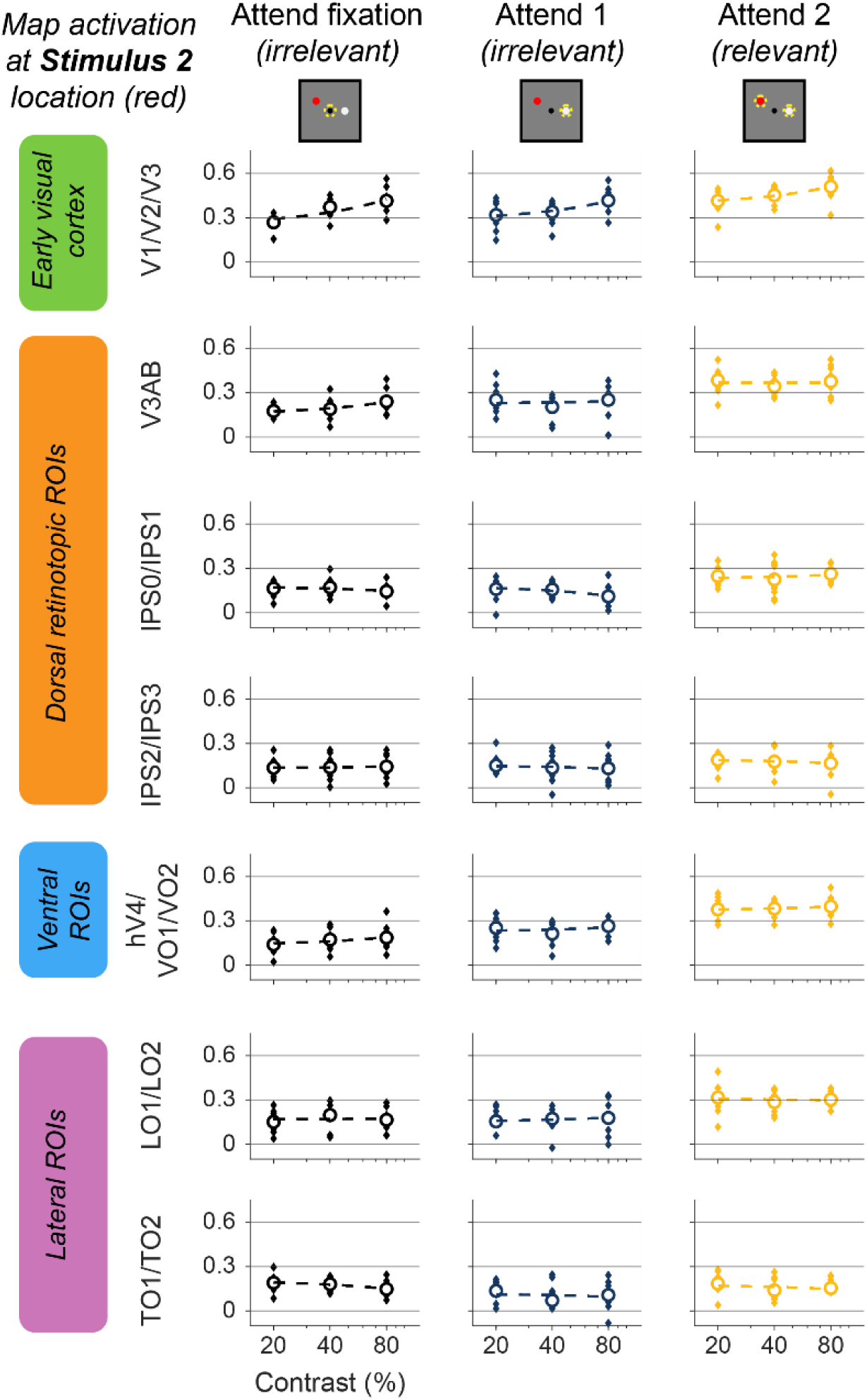
Map activation at Stimulus 2 location (related to Fig. 3). Data presented as Fig. 3D, with activation computed at the location of Stimulus 2. Stimulus 2 is task-irrelevant on Attend fixation trials and on Attend 1 trials, and is relevant on Attend 2 trials.

**Figure S3.**
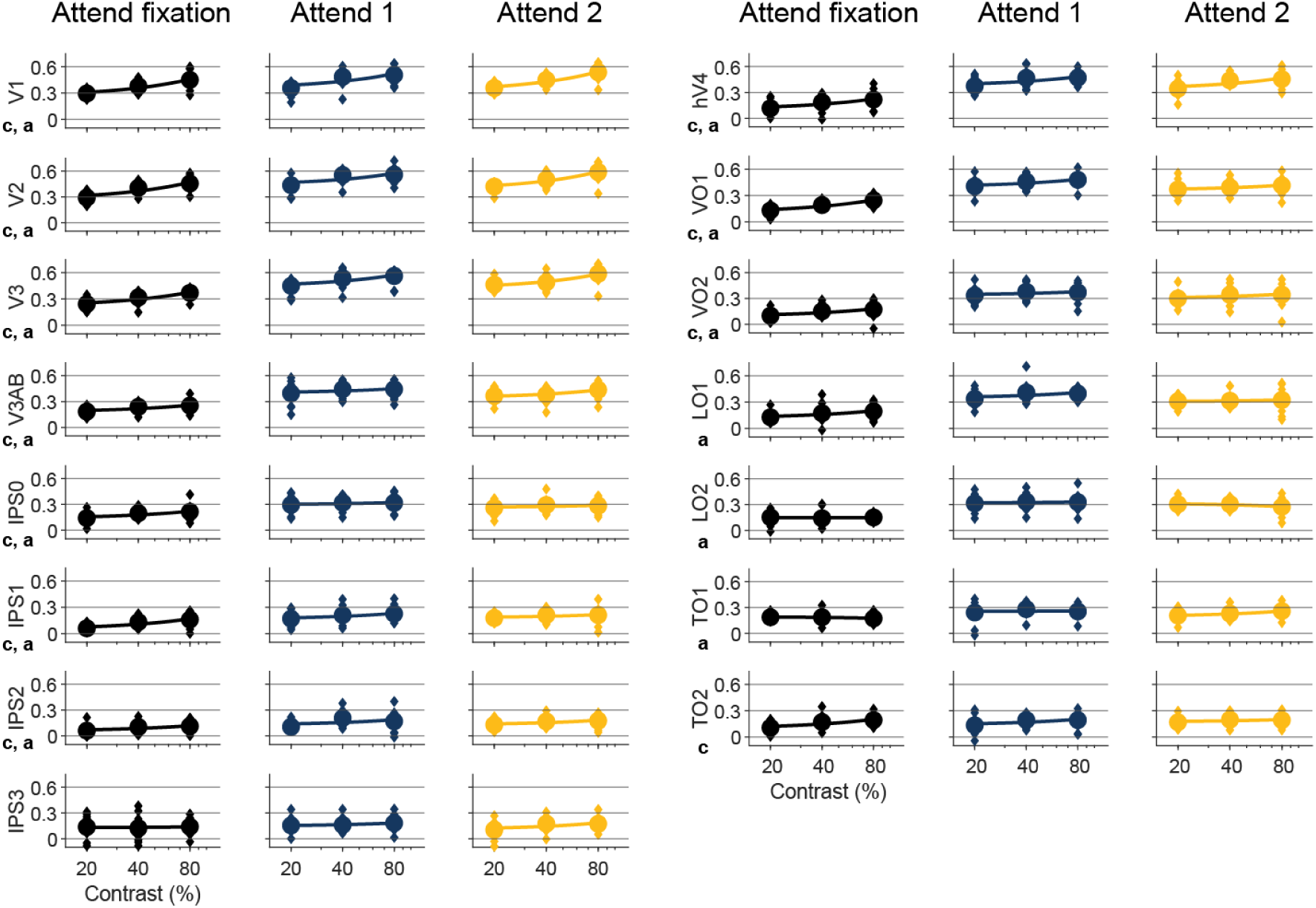
Contrast response functions across conditions for individual ROIs (related to Fig. 3). Data presented as in Fig. 3D, based on reconstructions computed from individual retinotopic ROIs. Statistics presented in Table S2.

**Figure S4.**
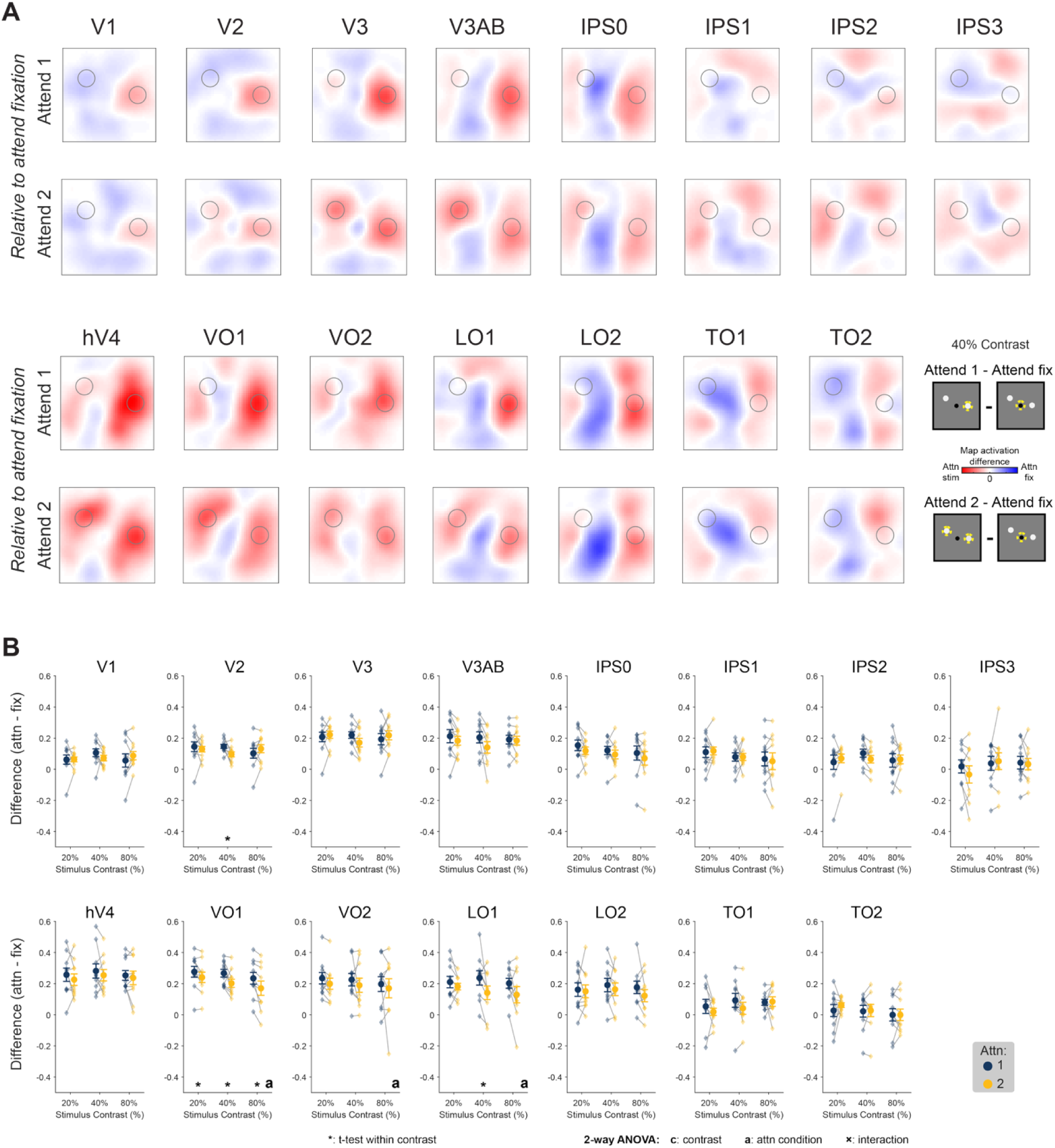
Relevance maps for individual retinotopic ROIs (related to Fig. 4). **(A)** Difference maps at 40% contrast between Attend 1 – Attend fixation (top) and Attend 2 – attend fixation (bottom). Data presented as in Fig. 4A. **(B)** Modulation of map activation at Stimulus 1 location at each contrast and for each task-relevance condition compared to Attend fixation. Data presented as in Fig. 4B. Statistics available in Table S4.

**Table S1.** *p*-values for effects of task relevance condition, stimulus contrast, and their interaction on map activation at the target stimulus location (related to Fig. 3)

|  | V1V2V3 | V3AB | IPSOIPS1 | IPS2IPS3 | hV4VO1VO2 | TO1TO2 | LO1LO2 |
| --- | --- | --- | --- | --- | --- | --- | --- |
| Condition | **<0.001** | **<0.001** | **<0.001** | **0.002** | **<0.001** | **0.038** | **<0.001** |
| Contrast | **<0.001** | **0.002** | **<0.001** | **0.008** | **<0.001** | **0.026** | 0.653 |
| Interaction | 0.459 | 0.525 | 0.728 | 0.501 | 0.856 | 0.606 | 0.455 |

To assess whether each region of interest (ROI) was sensitive to task relevance condition and stimulus contrast, we performed a 2-way repeated-measures ANOVA with within-subject factors of task relevance condition (Attend fixation, Attend 1, Attend 2) and matched stimulus contrast (20%, 40%, 80%), and their interaction. One ANOVA was performed per ROI. *p*-values are reported here; full statistics including *F*-values, degrees of freedom, and partial $\eta^2$ are available online (see *Data & code availability*). Bolded *p*-values indicate significant effects ( $p \leq 0.05$ ).

**Table S2.**
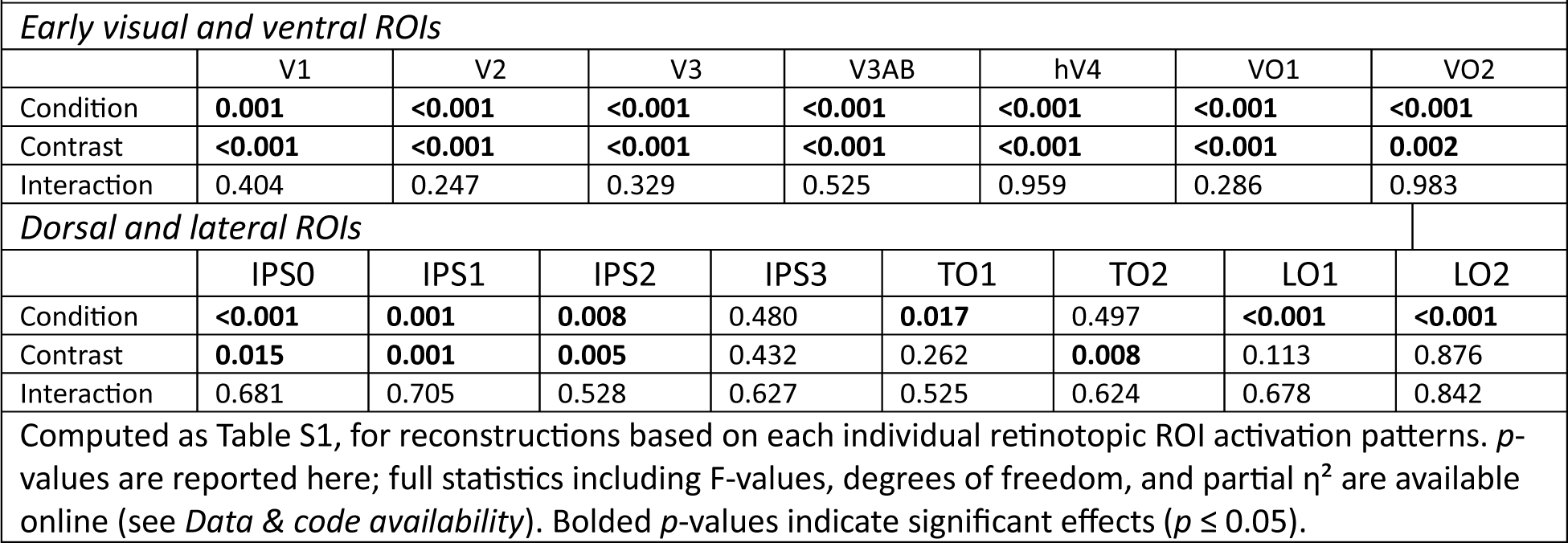
*p*-values for effects of task relevance condition, stimulus contrast, and their interaction on map activation at the target stimulus location for individual retinotopic ROIs (related to Fig. S3).

**Table S3.**
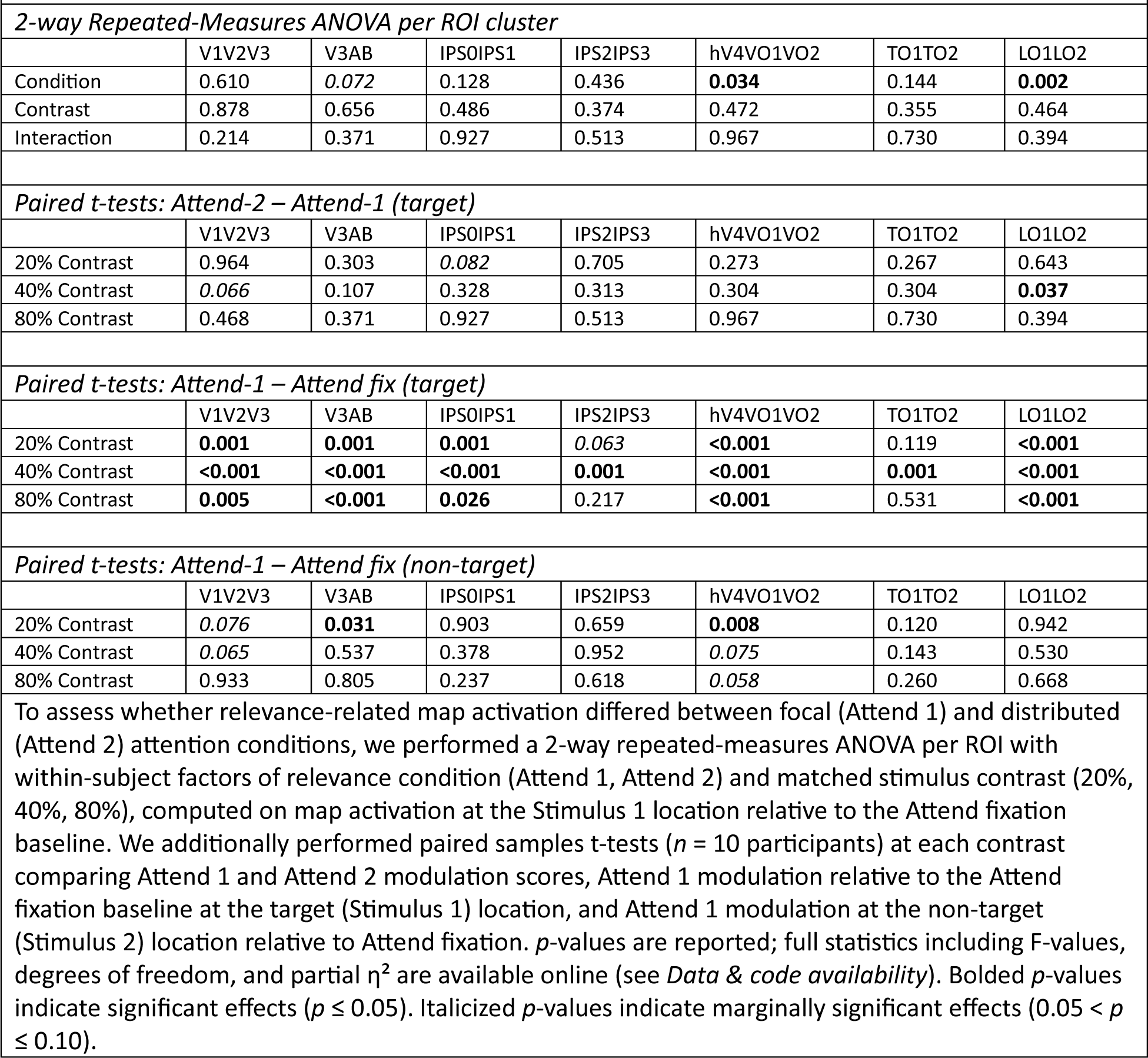
*p*-values for relevance-related modulations of map activation (related to Fig. 4).

**Table S4.** *p*-values for paired t-tests comparing relevance-related map modulation between Attend 1 and Attend 2 conditions at the target stimulus location for individual retinotopic ROIs (related to Fig. S4).

| Early visual and ventral ROIs |  |  |  |  |  |  |  |  |
| --- | --- | --- | --- | --- | --- | --- | --- | --- |
|  | V1 | V2 | V3 | V3AB | hV4 | VO1 | VO2 |  |
| 20% Contrast | 0.955 | 0.615 | 0.598 | 0.303 | 0.389 | **0.039** | 0.123 |  |
| 40% Contrast | 0.234 | **0.029** | *0.063* | 0.107 | 0.450 | **0.043** | 0.453 |  |
| 80% Contrast | 0.249 | 0.249 | 0.433 | 0.732 | 0.538 | **0.011** | 0.623 |  |
| Dorsal and lateral ROIs |  |  |  |  |  |  |  |  |
|  | IPS0 | IPS1 | IPS2 | IPS3 | TO1 | TO2 | LO1 | LO2 |
| 20% Contrast | 0.253 | 0.770 | 0.365 | 0.180 | 0.399 | 0.404 | 0.280 | 0.793 |
| 40% Contrast | 0.283 | 0.985 | 0.112 | 0.691 | 0.113 | 0.843 | **0.029** | 0.485 |
| 80% Contrast | 0.344 | 0.651 | 0.861 | 0.786 | 0.892 | 0.991 | *0.083* | 0.172 |
| Computed as Table S3, for reconstructions based on individual retinotopic ROI activation patterns. *p*-values are reported; full statistics including t-values and degrees of freedom are available online (see Data & code availability). Bolded *p*-values indicate significant effects ($p \leq 0.05$). Italicized *p*-values indicate marginally significant effects ($0.05 < p \leq 0.10$). | | | | | | | | |

